# Competitive hierarchies, antibiosis, and the distribution of bacterial life history traits in a microbiome*

**DOI:** 10.1101/2020.06.15.152272

**Authors:** Parris T Humphrey, Trang T Satterlee, Noah K Whiteman

**Affiliations:** Department of Organismic and Evolutionary Biology, Harvard University, Cambridge, MA. USA; Department of Ecology and Evolutionary Biology, University of Arizona, Tucson, AZ, USA; Department of Integrative Biology, University of California, Berkeley, CA. USA

**Keywords:** Pseudomonas, indirect interactions, phyllosphere, microbiome, phytopathogen

## Abstract

Microbiome manipulation requires an understanding of how species interact within communities. Can outcomes of ecological interactions be predicted from microbial life history traits, the identity of the species, or both? We addressed these questions by studying the competitive interaction network in a community of 40 endophytic *Pseudomonas* spp. bacterial isolates from a native plant. Pairwise competition experiments revealed competitive dominance of *P. fluorescens* over *P. syringae* strains within this microbiome-derived community. *P. syringae* strains with higher growth rates won more contests, while *P. fluorescens* strains with shorter lag times and lower growth rates won more contests. Adding to their competitive dominance, *P. fluorescens* strains often produced antibiotics to which few *P. syringae* strains were resistant. Many competitive outcomes among *P. syringae* strains were predicted to be reversed by *P. fluorescens* inhibitors because indirect benefits accrued to less competitive strains. *P. fluorescens* strains frequently changed competitive outcomes, suggesting a critical role of strains within this bacterial clade in structuring plant microbiome communities. Microbial traits also may provide a handle for directing the outcome of colonization processes within microbiomes.

## Introduction

The ecological forces shaping bacterial microbiome community structure are difficult to characterize, given the diversity and relatively uncultivable nature of these taxa, particularly in animals. Plants, in contrast, possess a highly cultivable microbiome and have potential to serve as models for understanding microbiome ecology and evolution generally. Moreover, plant growth-promoting bacterial (PGPB) formulations are being deployed in agriculture. Quantifying and predicting ecological outcomes among common species in these artificial communities is therefore also of practical value.

Competition may be the principle ecological force shaping microbial community structure (Foster & Bell 2012; Coyte & Rakoff-Nahoum 2019), yet distinct forms of competition can operate within communities: competition for shared resources and interference with another species’ ability to do so (Case & Gilpin 1974). In addition to structuring microbiome communities, competition of both types is a potent source of natural selection (Hibbing *et al.* 2010; Cornforth & Foster 2013; Mitri & Foster 2013). Teasing apart how exploitative and interference competition interact in a community context remains a challenge more generally (Amarasekare 2003; Delong & Vasseur 2013; Coyte *et al.* 2015). Furthermore, as diversity increases, the number of possible indirect interactions in the community scales faster than the number of direct interactions. Accordingly, a species may benefit from additional competitors if the net indirect effects dampen direct competition faced by other species (Levine 1976; Lawlor 1979; Stone & Roberts 1991; Wootton 1994; Miller & Travis 1996). Such indirect facilitation has not been well explored in microbiomes.

Species-rich communities are also more likely to harbor members with traits that have a large ecological impact (Banerjee *et al.* 2018). In microbial communities, strains that secrete diffusible antibiotics, resource substrates, or signaling molecules can alter the fitness of non-producers (Lee *et al.* 2010; Gutiérrez & Garrido 2019). By selecting for more specialized traits involved in resistance or metabolite uptake, these secretions can upend competitive hierarchies that would otherwise be mediated by canonical competitive fitness traits. It is unclear if microbial taxa with large indirect impacts are common in natural microbiomes (Banerjee *et al.* 2018). Leaf-dwelling (phyllosphere) bacteria secrete compounds altering growth and survival of nearby bacteria (Lindow & Brandl 2003; Quiñones *et al.* 2005; Dulla & Lindow 2009; Dulla *et al.* 2010) and can co-localize on the leaf surface and interior (Monier & Lindow 2005). Thus, there is potential for direct and indirect interactions between competing bacteria to affect community assembly and steady-state patterns of diversity in plant microbiomes.

Finally, competition need not be purely hierarchical: intransitive loops may arise in species-rich communities whereby numerical dominance cycles at local spatial scales, resulting in community stability (Kerr *et al.* 2002; Rojas-Echenique & Allesina 2011). Even modest intransitivity can buffer against extinction (Laird & Schamp 2006; Rojas-Echenique & Allesina 2011; Laird 2014) and the degree of intransitivity can shape species diversity (Reichenbach *et al.* 2007). Although intransitivity occurs in microbial systems in the laboratory (Kerr *et al.* 2002; Kelsic *et al.* 2015), its occurrence in natural microbiome communities is not well understood (Lankau *et al.* 2011; Godoy *et al.* 2017).

To address the various gaps highlighted above, we studied how microbial traits mediate direct and indirect competitive outcomes in an assemblage of co-occurring bacterial species from a wild, endophytic microbiome meta-community. Specifically, we (1) characterized life history trait variances and co-variances of diverse isolates in the laboratory, (2) examined how such traits related to competitive interaction networks manifest in spatial microcosms, and (3) analyzed whether indirect interactions among strains might be expected to strengthen or weaken competitive hierarchies among strains, with the latter expected to promote co-existence under natural conditions. We used a diverse set of endophytic *Pseudomonas* spp. bacteria derived from native bittercress (Brassicaceae: *Cardamine cordifolia* A. Gray), encompassing an extensive sample of the diversity found in both the putatively phytopathogenic *P. syringae* clade and the presumed saprophyte *P. fluorescens* clade (Humphrey *et al.* 2014; Humphrey & Whiteman 2020).

## Methods

### Overview

We measured a network of pairwise competitive interactions among 40 *Pseudomonas* spp. strains, wherein strains competed for shared resources in spatial microcosms. We quantified each strain’s ability to invade and defend against invasion and derived a composite measure of competitiveness that incorporated both invasive and defensive ability. We simultaneously measured each strain’s capacity to interfere with growth of surrounding competitors through inhibitory secretions, as well as each strain’s apparent ability to resist such inhibitors. Using independent measurements of maximum rate of increase, lag phase, and maximum yield *in vitro*, we then determined the underlying correlates of both exploitative and interference competitive abilities, as well as effect of phylogenetic distance on these correlations. Finally, using the distribution of pairwise outcomes measured in our competition assays, we inferred the number and direction of indirect interactions that would result in facilitation via inhibition of a superior competitor by a nearby producer strain.

### Bacterial strains

Of the 51 *Pseudomonas* spp. strains isolated from bittercress and previously described (Humphrey *et al.* 2014), we selected a set of 40 (26 *P. syringae*, 14 *P. fluorescens*) that represented the phylogenetic diversity present in this community (Humphrey & Whiteman 2020). We included the laboratory strain *P. syringae* pv. maculicola str. `ES4326` in our strain set owing to its phylogenetic similarity to strains isolated from bittercress and its extensive characterization in the laboratory as a pathogen of *Arabidopsis thaliana* (Cui *et al.* 2002, 2005; Groen *et al.* 2013). All bacterial strains used had undergone only one prior growth cycle after freezing following initial isolation on King’s B plates from surface-sterilized homogenates of bittercress leaf samples (Humphrey *et al.* 2014). For each strain, we estimated resource usage (i.e., growth) parameters (maximum growth rate *r*, lag phase *L*, maximum yield *K*) from *in vitro* growth cycles conducted in 96-well plates (see **Online Supplemental Materials [OSM]: Supplemental Methods** for details).

### Pairwise competition assays

We conducted pairwise high-density competition assays in spatial microcosms in which a “resident” strain inoculated onto the surface of each plate competed with each “invader” strain spotted on top (see **OSM: Supplemental Methods** for details). We visually scored growth of each invader as 0 for no visible growth of the invader above a negative control spot containing sterile growth media alone, 0.5 for a largely translucent ‘megacolony’, which reflected a definite presence of growth but which was relatively suppressed and confined to the megacolony margin, and 1 for obvious and robust megacolony growth. We scored inhibition interactions as a binary outcome indicating the presence of a zone of clearance (halo) ≥ 1 mm surrounding the extent of the invader megacolony.

### Calculating indexes of competitiveness

Each strain was assayed under 40 different conditions both as resident strain and invader, comprising an interaction network with 1,600 entries (including self vs. self). One version of the interaction network represents the outcome of resource competition and details the extent of growth of each invader, while the other captures the presence or absence of inhibitory interactions indicated by zones of clearance in the resident population. For resource competitions, we calculate the invasive capacity (*C*_*o*_) and defense capacity (i.e. territoriality; *C*_*d*_) of each strain. *C*_*o*_ for each strain *i* was calculated as

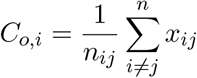

where *x*_*ij*_ ∈ {0, 0.5, 1} and *n*_*ij*_ is the total number of scored interactions for each strain as the invader with all non-self resident strains. *C*_*o*_ is thus the expected value of growth attained by each strain as the invader across the population of residents. Similarly, *C*_*d*_ quantifies the ability of each strain to resist invasion by other strains and is calculated as

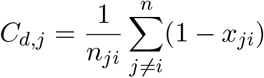

Here, strain *j* is in the resident state, and *x*_*ji*_ ∈ {0, 0.5, 1} as before but with a subscript reversal, indicating the degree to which the resident prevented the growth of each invader *i*. As above, *n*_*ji*_ is the number of interactions occurring between each focal resident and its non-self invaders. *C*_*d*_ can thus be interpreted as the expected amount of growth each resident strain can prevent among the population of invaders assayed.

We then calculated an overall exploitative competition index, *C*_*w*_, for each strain as

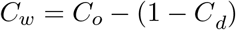

where −1 ≤ *C*_*w*_ ≤ 1. These extremes represent absolute competitive inferiority (−1), where a strain failed to prevent all growth of any invader and similarly failed to invade any other strain, to absolute competitive dominance (1), where a strain fully invaded all residents and fully prevented growth of all invaders.

We also calculated *C*_*t*_ and *C*_*r*_ based on the interaction matrix for interference competition. Here, *C*_*t*_ is the proportion of successful invasions (i.e., given growth of 0.5 or above) that also resulted in halo formation produced by the invading strain, indicating inhibition of the resident. *C*_*r*_ for a strain is the proportion of contests with all invading inhibitor strains (i.e., all strains with *C_t_* > 0) that failed to result in halo formation, which we interpreted as resistance. Analogous to *C*_*w*_ above, we calculated an overall interference competition index, *I*_*w*_, as

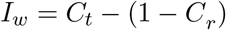

where −1 ≤ *I*_*w*_ ≤ 1, which is analogous to the aggressiveness index (*AI*) of (Vetsigian *et al.* 2011).

### Analyzing the distribution of competitive outcomes

We determined when outcomes of all pairwise interactions between strains *i* and *j* (*i* ≠ *j*) took the following forms: reciprocal invasibility (`RI`), where strains *i* and *j* each invade one another; reciprocal non-invasibility (`RNI`), where strains *i* and *j* cannot invade each other; and asymmetric (`Asym`), where strain *i* invades strain *j* but *j* cannot invade *i*. To compare outcome distributions, we constructed binomial linear models to estimate the probability of `RI, RNI`, and `Asym` as a function of bacterial clade (*P. syringae* versus *P. fluorescens*).

In addition, we compared trait co-variances and overall levels of trait dispersion between *P. syringae* and *P. fluorescens*, correcting for phylogenetic distance between strains in each clade. To do so, we first we conducted principal components analysis (PCA) using the matrix of mean-centered and scaled competitive indexes and growth parameters for all strains (40 × 9 matrix) as input. We then calculated Euclidean distance between vectors of [`PC1, PC2, PC3`] for all pairs of strains within each *Pseudomonas* clade. Using these calculated pairwise multivariate trait distances as a response variable, we computed linear regression models with bacterial clade as well as phylogenetic distance (*D*_*g*_) as predictors. We calculated *D*_*g*_ as the pairwise uncorrected nucleotide distance between 2,690 bp of sequence comprised of four partial housekeeping gene sequences previously generated for each strain from Humphrey et al. (2014). Orthologous sequences from the genome of Psm4326 were derived from its published genome sequence (Baltrus *et al.* 2011); RefSeq ID `NZ_AEAK00000000.1`).

### Inferring indirect interactions from the pairwise network

We next examined the structure of the pairwise competitive interaction network for signatures of intransitivity (i.e., non-hierarchical or context-dependent interactions). Using data from pairwise interaction outcomes, we assessed (1) whether three-strain competitions would result in intransitive loops (e.g., rock-paper-scissors outcomes) such that no species would be globally dominant; and (2) whether the presence of secretions from a nearby *P. fluorescens* strain would reverse the outcome of a pairwise interaction that would typically result in competitive dominance of a single strain (indirect facilitation). Facilitation can occur by strain `A` releasing strain `C` from inhibition from strain `B` (where `A` also has to be resistant to `B`’s inhibitors), or from resource competition from superior competitor strain `B`. This analysis is agnostic to mechanism but calculates the proportion of conditions under which facilitation of an otherwise weaker competitor is expected to arise. A total of 8,203 trios were analyzed for potential facilitation based on the pairwise interaction data from 641 pairs of strains that met the competitive asymmetry criteria.

For each strain, we calculated the net effect of antagonistic vs. facilitative indirect interactions across all possible trios and compared this to underlying fitness metrics derived from the pair-wise interaction network. We then compared how strongly the net effects from indirect facilitation are expected to change fitness ranks of strains in relation to their baseline values of overall competitiveness (*C*_*w*_).

## Results

### Competitive outcomes

Pairwise soft-agar invasion assays revealed that the competitive ability of *P. fluorescens* strains was consistently superior to *P. syringae* strains (Fig. 1): ~99% of strain pairings between the two clades resulted in asymmetric dominance of *P. fluorescens* over *P. syringae* (99% `Asym`; Fig. S2; Tables S1, S2). Within *P. fluorescens*, the proportion of reciprocally non-invasible (`RNI`) pairings was significantly higher compared to within *P. syringae* pairings (Fig. S2; Tables S2, S3). The competitive dominance of *P. fluorescens* over *P. syringae* was evident across both exploitative and interference-based measures of competitiveness (Figs 1, 2; Table S1).

**Figure 1:**
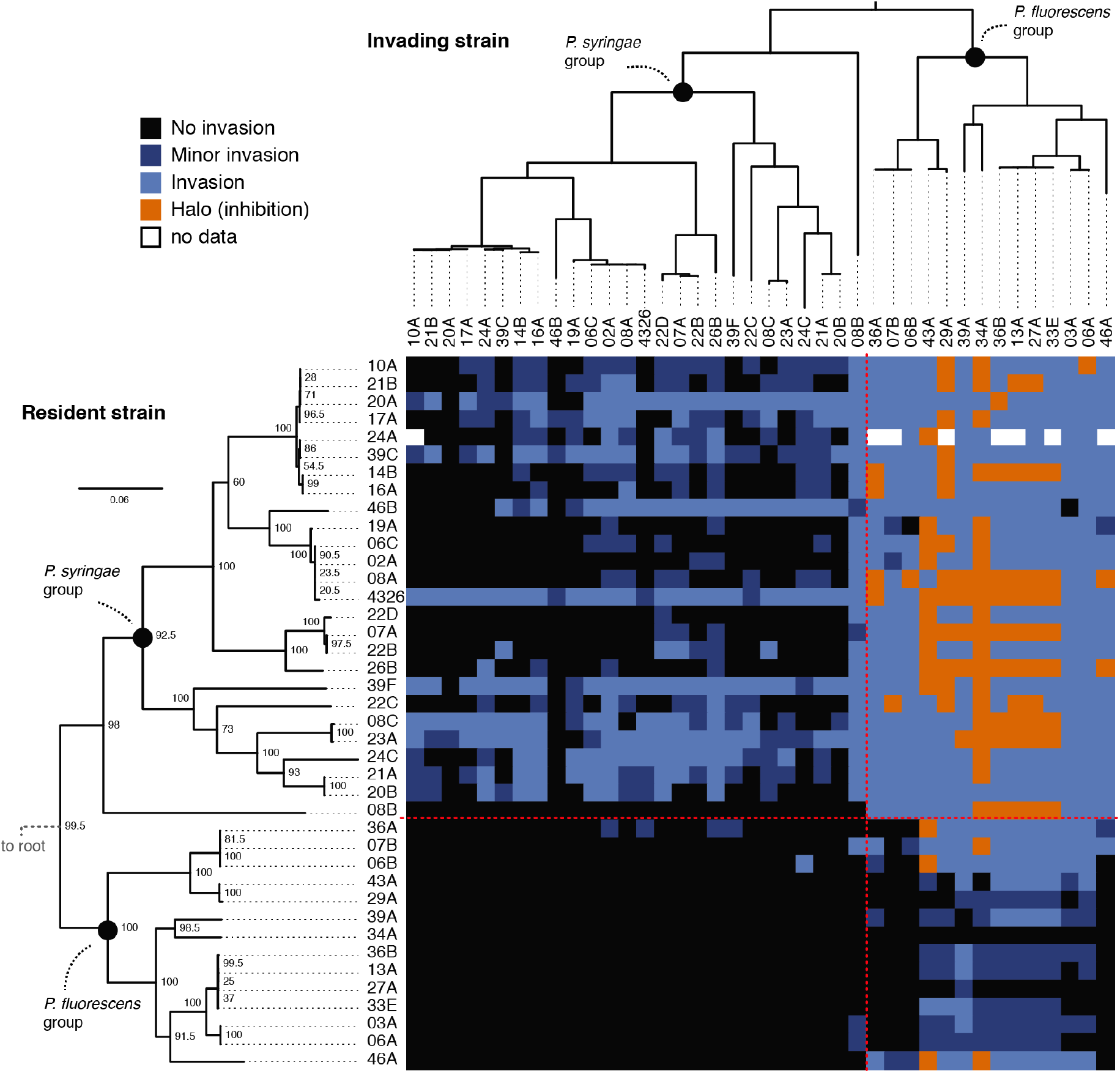
Pairwise competitive interactions in a phyllosphere *Pseudomonas* spp. community. Rows reflect strains in the resident state, while columns reflect strains in the invader state. Dashed red lines through interaction matrix denote within–between clade divisions for ease of visualization. Phylogeny modified from Humphrey et al. (2014).

**Figure 2:**
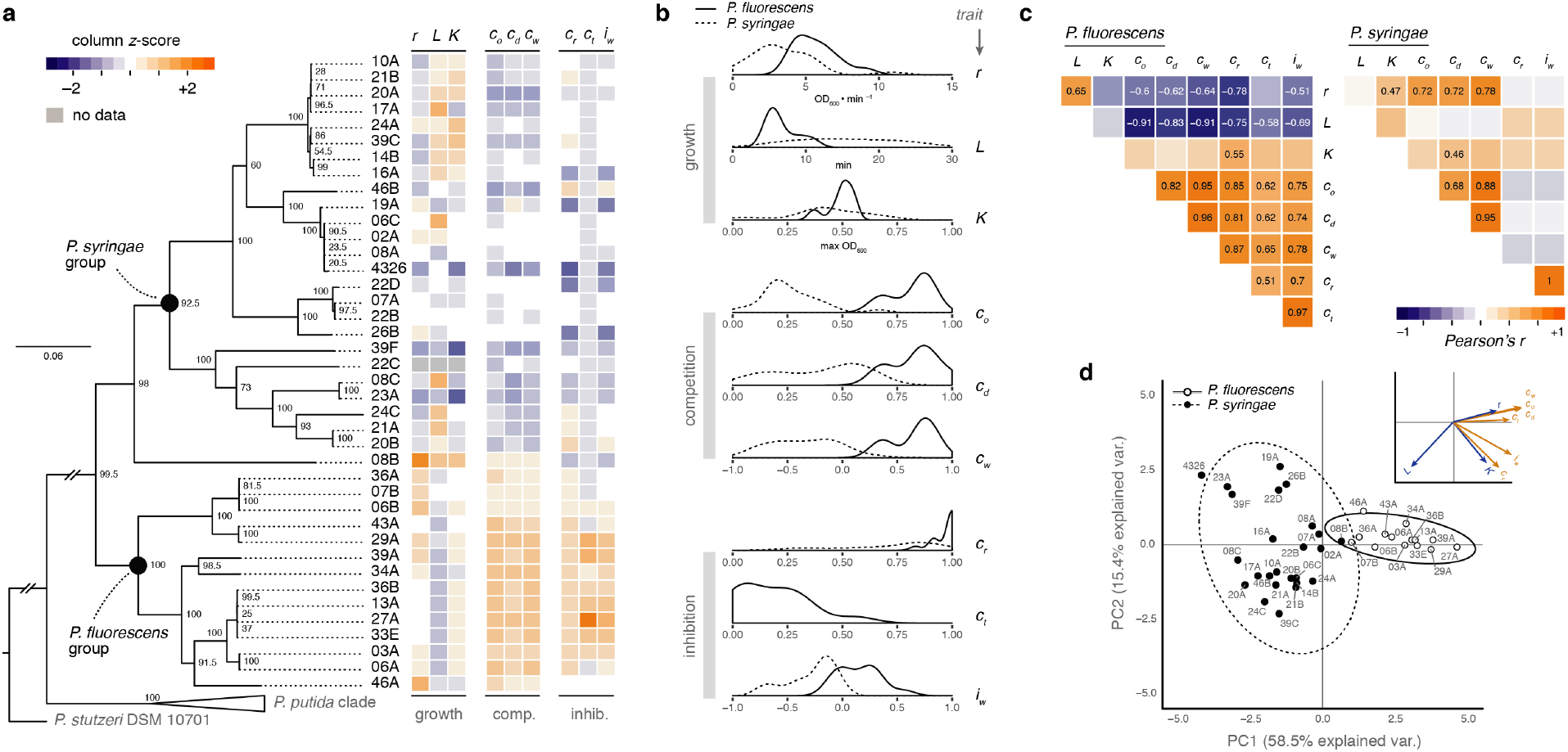
Phylogenetic distribution of life history trait variation within a *Pseudomonas* spp. community. **a**. Life history components are maximum growth rate (*r*_*m*_), lag phase (*l*), maximum yield (*K*), derived from individual microcosm growth experiments; and components of offensive (*C*_*o*_), defensive (*C*_*d*_), overall (*C*_*w*_) competitiveness, resistance to toxicity (*C*_*r*_), toxicity (*C*_*t*_), and overall interference capacity (*I*_*w*_) derived from a pairwise competitive interaction network (see *Methods*). Column *z*-score of each trait value indicated by color. **b**. Smoothed frequency distributions of trait values for each measured trait by clade (*P. fluorescens* and *P. syringae*). **c**. Pairwise correlations and principle component analysis (PCA) (**d**) of nine traits revealed clear dissimilarities in trait distributions and patterns of co-variance between clades. Correlations with text values reflect magnitude of each Pearson’s *ρ* where the FDR corrected *p* < 0.05; comparisons with FDR-corrected *p* < 0.10 are italicized. **d**. PCA 95% envelopes per clade depicted as solid or dashed ellipses. Dots are labeled with strain IDs. Individual trait vector loadings are in blue for resource use traits and orange for competitive indexes).

### Interference competition

Of the 40 strains assayed, 13 (all *P. fluorescens*) produced halos surrounding some subset of the resident strains they invaded (antibiosis), indicating the production of antibiotics (diffusible inhibitors/toxins) (Fig. 1). Mean inhibition index (*I*_*w*_) among *P. fluorescens* strains was 0.15, although two strains inhibited only one other, and *P. fluorescens* strain `03A` failed to inhibit any strain (Fig. 1). Four *P. fluorescens* strains were susceptible to inhibition by two of the toxic strains (`43A`, `34A`; Fig. 1). Resistance to toxin producers in *P. syringae* was variable, although the mean value was high at 0.72 (Fig. 2b; Table S1).

In at least one case, resistance among *P. syringae* strains showed a strong correlation with phylogenetic position: invading strain *P. fluorescens* str. `43A` adopted distinctly different phenotypes in pairings with *P. syringae* strains from different sub-clades (`perMANOVA` *F* = 7.04, 1000 permutations, *p* = 0.002; Fig. S3). Nine of the 25 `43A` megacolonies had a smooth morphology, 13 adopted a highly motile morphology we call the “smooth spreader”, and the three remaining adopted a wrinkly spreader-like morphology (Fig. S3a-c). Inhibitor production by `43A` was strongly associated with the smooth morph (*χ*^2^ = 19.2, *p* < 0.001; Fig. S3e); `43A` only inhibited one strain as the smooth spreader morph, and then only after it had stopped expanding across the plate (personal observation). None of the three wrinkly spreader-like morphs produced toxins that inhibited a resident strain.

### Life history correlates of competitiveness

The correlations between competition and growth traits showed opposite patterns for strains within *P. syringae* versus *P. fluorescens*: overall exploitative competitiveness (*C*_*w*_) was negatively correlated with both *r* and *L* for *P. fluorescens* (Pearson’s *ρ* = −0.78, −0.75, respectively; Fig. 2c). That is, *P. fluorescens* strains with shorter lag (smaller *L*), and thus smaller *r*, were more competitive in our assay. This apparent trade-off between maximum *in vitro* growth rate *r* and growth initiation (1*/L*) was not observed across *P. syrinage* strains. Instead, *C*_*w*_ in *P. syringae* was positively correlated with only *r* (*ρ* = 0.78; Fig. 2c). Strains from neither clade showed a canonical trade-off between *r* and *in vitro* saturation density (*K*). On the contrary, *P. syringae* strains showed a positive correlation between *K* and growth rate as well as defensive capacity *C*_*d*_, while for *P. fluorescens K* was positively correlated with levels of resistance (*C*_*r*_). Overall, offense (*C*_*o*_) and defense (*C*_*d*_) were strongly positively correlated with linear slopes near 1 for both clades (Fig. 2c; Fig. S5). All three measures of exploitative competition were positively related to interference measures for *P. fluorescens* (Fig. 2c).

Principal component analysis (PCA) of all nine traits revealed largely non-overlapping 95% confidence ellipses for the two clades (Fig. 2d). The first two PCs together explained 72.5% of the variation in the data. The loading vectors of *C*_*w*_ and lag duration were in opposing directions, indicating a negative correlation, while those for competitiveness and inhibitory capacity are largely co-linear, indicating a positive correlation (Fig. 2d). The loading for resistance, *C*_*r*_, was nearly co-linear with lag duration, a relationship not apparent in the pair-wise correlation analysis in Fig. 2c.

Overall, strains within the *P. syringae* clade showed greater intra-clade pairwise trait differences across PCs 1-3 than strains within *P. fluorescens* (Welch’s unequal variants *t* test, *t* = 8.7, *p* < 10^−6^; Fig. S7). While multivariate trait distance increased on average with phylogenetic distance (*D*_*g*_ term *β* = 0.1, *p* < 10^−10^; Table S4), *P. syringae* strains showed a higher average trait distance even after accounting for *D*_*g*_ in a multiple regression model (`Psyr` term *β* = 0.9, *p* < 10^−8^; Table S4).

### Competitive interaction network and intransitivity

Five trios met the criteria for a rock–paper–scissor game out of the 9,604 possible trios of interactions evaluated (Fig. 3a). Nine unique strains were implicated in these trios. Each trio was comprised of distantly related *P. syringae* strains (mean *D*_*G*_ between strains in R–P–S trios = 0.118 [0.115–0.122 95% CI]). A further 632 (7.7%) met the criteria whereby the inferior competitor was facilitated by the inhibition of the superior competitor by a third party to which the facilitated strain was resistant (Fig. 3a).

**Figure 3:**
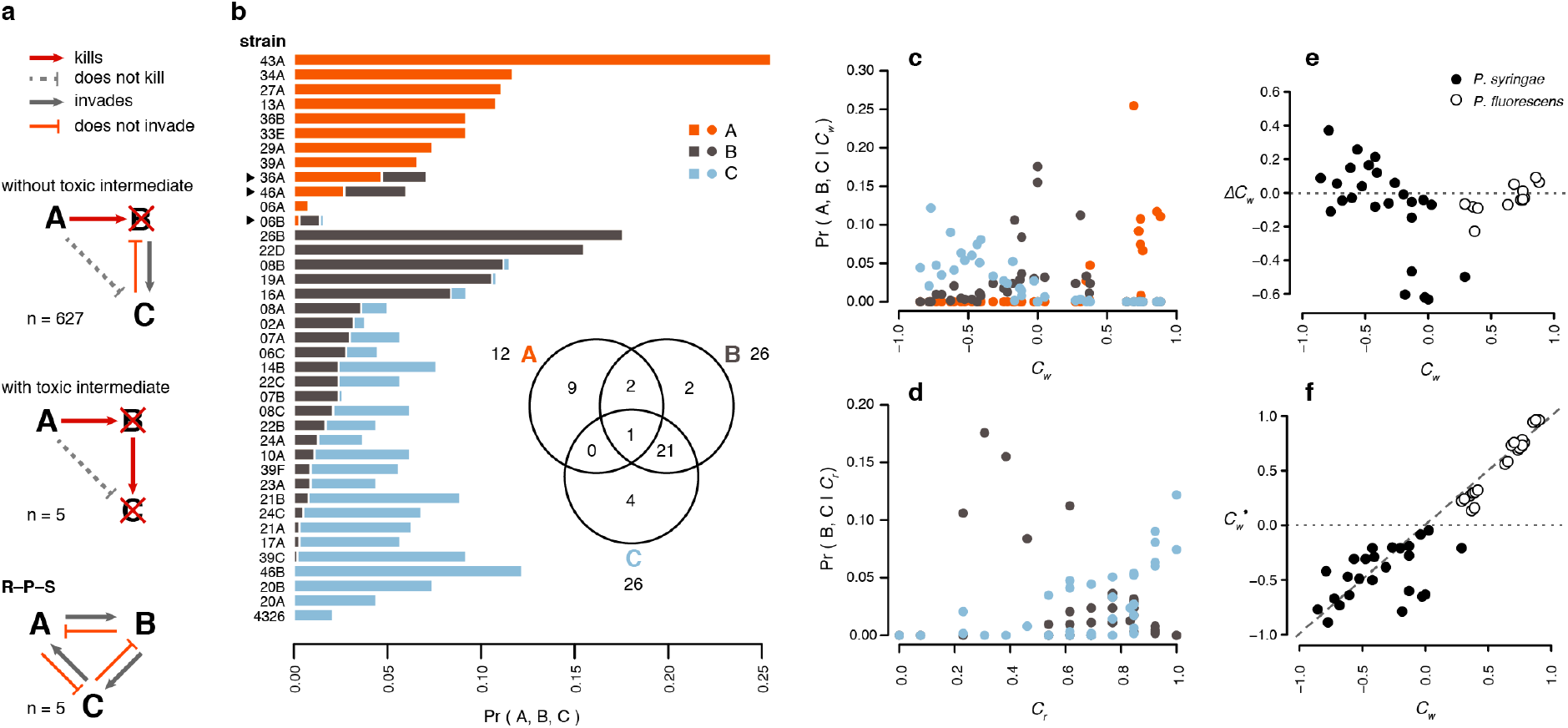
Prevalence of intransitive interactions in a *Pseudomonas* spp. interaction network. **a**. Types of interaction trios resulting in facilitation or rock-paper-scissors (R–P–S) competitive asymmetries. *N* = number of trios meeting the given criteria out of the total trios analyzed (see Methods). **b**. Frequency distributions of how often each strain played the facilitator (*A*), the knocked-out intermediate (*B*), or the facilitated (*C*). Several strains played multiple roles; strains in facilitative trios with as well as without toxic intermediates are indicated with black triangles to the left of the strain IDs. Panel (**b**) inset displays the distribution of the number of unique strains that played each combination of roles. Only `06B` played all three roles. The probability of playing *A*, *B*, or *C* roles in facilitative trios varied with (**c**) overall competitiveness, *C*_*w*_, as well as (**d**) resistance. **e.** Δ*C*_*w*_ plotted against base-line *C*_*w*_ shows initially weaker *P. syringae* strains benefit the most from indirect interactions (Pearson’s *ρ* = −0.67), while *P. fluorescens* fitness remains relatively unaffected by indirect interactions (*ρ* = 0.74). **f.** Net competitive fitness 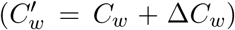 after considering indirect effects weakens competitive hierarchies among *P. syringae* (*ρ* = 0.50) but has little effect on *P. fluorescens* competitive fitness (*ρ* = 0.98).

Despite the overall tendency to reinforce pairwise interactions, indirect facilitation from inhibitor-producing strains implicated nearly all (39) of the 40 studied strains in one or more of the three possible trio roles: the facilitator, the knocked-out competitor, or the facilitated strain (*A*, *B*, and *C*, respectively; Fig. 3a). Overall, 26 strains were facilitated (*C*), and 21 of these also served as the knocked-out competitor (*B*) in a subset of the trios (Fig 3b, inset). Twelve of the 13 inhibitor-producing strains (all *P. fluorescens*) were implicated as facilitators (*A* strains) (Fig 3b, inset).

Intuitively, the propensity towards *B* vs. *C* roles was correlated by underlying differences in competitive fitness: the most facilitated strains (high *C* fraction) were among the least competitive (low *C*_*w*_) in the population, indicated by a negative correlation (*r* = −0.76 [|0.86 − 0.58| 95% CI], *p* < 10^−5^; Fig. 2c). *B* strains were intermediate relative to the entire range of *C*_*w*_ values. Facilitator *A* strains had consistently higher *C*_*w*_, owing to the generally higher competitiveness of *P. fluorescens* strains: in all but 6 of the 632 facilitation trios, the *A* strain out-competed the *C* strain in the pairwise network, even though such strains were resistant to their inhibitors (Fig 3b). This finding suggests that facilitation in this network depends on it occurring at a distance whereby the facilitator does not immediately out-compete the resistant strain which it facilitated. Also intuitively, resistance (*C*_*r*_) was strongly positively correlated with the probability of being facilitated (Pearson’s *ρ* = 0.57 [0.32 – 0.75 95% CI], *p* < 10^−4^.

Only rarely were *P. fluorescens* strains anything other than the facilitator strain: only three were ever knocked out by an *A* strain to which they lacked resistance (`36A, 46A, 06B`). This finding reveals that *P. fluorescens* strains very rarely benefit from indirect facilitation, in contrast to their frequent role as facilitator (Fig. S8). One strain (*P. fluorescens* str. `43A`) played the role of facilitator (*A*) in >25% of all facilitation trios, over 2.5-fold more often than the next most frequent facilitator (Fig. 3b). This indicates that the presence of individual inhibitor-producing community member can substantially shift the outcome distribution among non-producers.

Averaged across all inhibitor-producing strains, the net effects of indirect interactions reshuffled the fitness ranks of *P. syringae* strains to a degree that weakens the original pairwise competitive hierarchy (rank correlation *ρ* between *C*_*w*_ and 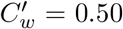; Fig. 3e; Fig. S8), such that the former advantage of several top *P. syringae* competitors gets redistributed across a larger number of relatively weaker competitors. In contrast, the hierarchy among *P. fluorescens* strains was generally recapitulated, or exacerbated, by indirect interactions arising from antibiosis in this network.

## Discussion

### Overview

We discovered clear clade- and trait-level associations with the outcomes of competitive interactions among naturally occurring bacterial strains. Using a subset of endophytic bacteria isolated from a native sub-alpine plant (*C. cordifolia*), we found major differences in both exploitative and interference competitiveness between the two principle *Pseudomonas* spp. clades in this endophytic community. Trait co-variance structure revealed the biological differences between these two major clades of native plant-associated *Pseudomonas* spp. bacteria. Such patterns suggest that the evolution of competitiveness may involve distinct components of life history in these bacterial lineages. When placed into an ecological context, the trait distributions we revealed across this bacterial assemblage are predicted to generate context dependence in competitive outcomes in the form of facilitation, whereby a inhibitor strain displaces a strong competitor and thereby facilitates a resistant but weaker recipient. Thus, the community context of interference competition is important for predicting the outcome of competitive pairings which typically depend primarily on exploitative capacity.

Such a dataset allows dissection of several dimensions of *in vitro* fitness exhibited by a natural community of phyllosphere *Pseudomonas* spp. and provides a platform for testing hypotheses about the mechanistic bases of competitive traits (e.g., toxin production and resistance) and their potential effects on ecological diversity and microbiome community structure. We also showed that *P. fluorescens*, presumed to be soil dweller, can be both common and important in structuring the outcome of ecological interactions within the context of the leaf microbiome. Together, this work helps build an understanding of how competitive traits might evolve in tandem with other life history traits in representatives from real communities that interact in nature.

### Correlations between growth traits and competitiveness

Neither *P. syringae* nor *P. fluorescens* strains exhibited canonical growth rate trade-offs with maximum yield, *K*, which can result in a tragedy of the commons whereby rapid but wasteful use of resources yields higher competitive ability (Pfeiffer *et al.* 2001; MacLean 2008). Rather, a more pronounced signal was that maximum growth rate was correlated with a longer lag phase in *P. fluorescens*. This pattern contradicts the traditional dichotomy between generally “fast” vs. “slow” life histories and contrasts with patterns observed in microbial evolution experiments. For example, *Escherichia coli* lines adapting to a glucose-limited environment exhibited coordinated increases in growth rate and shorter lag time after 10,000 generations (Vasi *et al.* 1994; Lenski *et al.* 1998). Additionally, *E. coli* selected to persist in lag phase during periods of antibiotic stress exhibited no pleiotropic changes in maximum growth rate despite up to a 10-fold increase in lag time (Fridman *et al.* 2014). Our study adds support for the idea that lag phase deserves attention as an important feature of microbial life cycles, and characterizing the physiology of cells during this phase may reveal the nature of its correlations with maximum growth rate and competitive fitness in this and other systems.

The negative correlation between lag phase and growth rate in *P. fluorescens* resembles a colonization-competition trade-off. Spatial priority effects arising from territoriality can provide a mechanism for maintenance of colonization–competition trade-offs that would otherwise lead to competitive exclusion (Edwards & Schreiber 2010). A colonization–competition trade-off underlies territoriality in *Vibrio* spp. based on the differential ability of clones to contest territory vs. disperse to new ephemeral habitats (Yawata *et al.* 2014). One hypothesis arising from our work is that *P. fluorescens* strains that preempt as much space as possible within patchy and ephemeral leaf environments may reap the rewards of their territorial monopoly even at the expense of a decreased maximal growth rate.

The production of exudate (*C*_*t*_) or exudate resistance (*C*_*r*_) did not trade-off with any of the life history traits we measured (Fig. 2a). This is consistent with findings that exudate production did not affect *in vitro* growth rates measures in *P. fluorescens* (Garbeva *et al.* 2011). Instead, we found a positive correlation between inhibitory ability (*C*_*t*_) and overall exploitative competitiveness for *P. fluorescens*. Although perhaps unexpected from a theoretical perspective (Neumann & Jetschke 2010), such a positive correlation is nevertheless intuitive: megacolonies invading a resident strain presumably must reach a critical size in order for any toxicity to be detectable if induction is either density dependent or if the toxic effects are concentration dependent. Cells may only reach such a critical density if their relative exploitative competitiveness enables them to do so, without which interference competitive ability is irrelevant. Further empirical work, scaling from individual cells to populations, will be required to properly ground co-existence theory for microbes in mechanistic models of trait-trait interactions.

Our study is limited in that we relied on visible manifestation of growth inhibition. Interference mechanisms range from direct injection of bacterial effectors via Type VI Secretion Systems (Decoin *et al.* 2014, 2015), the production of subversive growth-regulating secreted N-acylhomoserine lactones (AHLs) or enzymes that quench these signals typically involved in quorum sensing (Dulla & Lindow 2009; Dulla *et al.* 2010), or the production of secreted toxins (e.g. bacteriocins or phage-derived proteins). Further work is needed to describe the range of interference mechanisms that may operate within plant microbiomes and to characterize the ecological effects of newly described modes of interference capable of being deployed by *P. syringae* (Hockett *et al.* 2015; Kandel *et al.* 2020) that this study was not capable of detecting.

### Ecological implications

If strains from *P. syringae* and *P. fluorescens* were to compete in an unstructured environment, where preemption of space was irrelevant, *P. syringae* strains with high growth rates might be expected to out compete a variety of *P. fluorescens* strains with relatively lower growth rates (Fig. 2). But within the structured and ephemeral context of the leaf environment, *P. fluorescens* may act as a territorial species whose potential effect in the phyllosphere may be to exclude colonization by other strains including *P. syringae*. This is consistent with the identity of *P. fluorescens* as a plant mutualist, although the evidence of this comes exclusively, to our knowledge, from studies of its indirect effects via plant defensive signaling or direct toxicity to pathogenic fungi following its colonizing of plant roots (Mendes *et al.* 2011; Hol *et al.* 2013). In addition to such indirect effects, the superior competitiveness of *P. fluorescens* over *P. syringae* suggests that direct interactions may affect phyllosphere bacterial community assembly and plant disease risk from phytopathogenic isolates of *P. syringae*. Irrespective of the underlying mechanisms of interference and resistance, the frequency of these traits in a community may have large indirect effects that generate context-dependent competitive asymmetries among diverse genotypes.

The ecological context in which traits are expressed impacts functional diversity (both genetic and phenotypic) found within natural communities (Ohgushi *et al.* 2012), despite strong pairwise competitive asymmetries, as seen here between *Pseudomonas* spp. clades. In our interaction network, indirect effects of interference competition may equalize fitness differences between *P. syringae* competitors that otherwise have asymmetric exploitative abilities (Fig. 3b; Fig. S8). Facilitation of the sort explored here is only possible with an intermediate frequency of toxin resistance expressed by *P. syringae* (Fig. 3d). The fact that resistance is not more common among *P. syringae* suggests a cost of resistance that did not manifest itself in the assays conducted in our study. Further study into the mechanisms of production of, and resistance to, interference traits in this community would help explain the distribution of these traits in the community as well as their costs and correlations with other traits.

We show that the gains from facilitation are predominantly accrued by weaker resource competitors (Fig. 3c-f; Fig. S8). Only in a small subset of the facilitation trios could the facilitated strain invade the producer. When the facilitated strain does not pose a competitive threat to the facilitator—as is the case most of the time here—the gains from facilitation may be short-lived. However, the overall effect of this degree of facilitation may be to prolong periods between exclusion/extinction events, elevating the diversity that is observable at any given point within the system (Laird & Schamp 2006). The additional form of intransitivity found in our study is a pair of extended trios that have R–P–S invasion asymmetries, which are predicted to lead to frequency-dependent or cyclical invasion dynamics (Laird 2014). This prediction is awaiting an empirical test, and this system presents an excellent opportunity for doing so.

### Conclusions

We found that competitive abilities of strains within a natural assemblage of plant-derived *Pseudomonas* spp. varied between the two major clades present, *P. fluorescens* and *P. syringae*. Competitive fitness in our assays hinged on different traits in these two clades, and the higher degree of inter-strain trait dispersion in *P. syringae* may indicate that the focal traits measured here undergo more rapid evolution given the same degree of phylogenetic divergence (Fig. 2d; Fig. S7). We found no apparent life history trade-offs between growth rate and yield. Although speculative, the *P. fluorescens* clade may contain early colonizing strains that contest territory to a greater extent, which may serve to directly buffer against leaf colonization from potentially phytopathogenic *P. syringae*. In contrast, a high degree of inhibitor resistance among *P. syringae* may prevent local exclusion when spatial structure releases them from direct exploitative competition with *P. fluorescens*. Finally, the combination of exploitative and interference competition due to inhibitor-mediated facilitation may stabilize co-existence of strains that otherwise competitively exclude one another. Our study sheds light on the types of ecological interactions between bacterial lineages within microbiomes that should be quantified during development of microbial formations for clinical and crop enhancing purposes.

## Acknowledgements

P.T.H. and N.K.W. gratefully acknowledge funding from the National Science Foundation (Grant Nos. DEB-1309493 to P.T.H. and DEB-1256758 to N.K.W.), and the National Institute of General Medical Sciences of the National Institutes of Health (Grant No. R35GM119816 to N.K.W.).

