## Supplementary material for "Competitive hierarchies, antibiosis, and the distribution of bacterial life history traits in a microbiome*": Online Supplemental Material

**Parris T Humphrey**

**Trang T Satterlee**

*Department of Ecology and Evolutionary Biology, University of Arizona, Tucson, AZ, USA*

**Noah K Whiteman**

*Department of Integrative Biology, University of California, Berkeley, CA. USA*
*Department of Ecology and Evolutionary Biology, University of Arizona, Tucson, AZ, USA*

#### Supplemental Methods

##### *In vitro* growth conditions

We characterized canonical growth traits of each bacterial strain separately by tracking cell density through a complete lag–exponential–saturation cycle *in vitro*. We re-streaked each strain from -80 C stocks onto King's B (KB) agar plates +10 mM MgSO4 and incubated at 28 C for 3 days. We picked and inoculated single colonies into 1 mL minimal media at pH 5.6 ('MM'; 10 mM fructose, 10 mM mannitol, 50 mM KPO4, 7.6 mM (NH4)2SO4, and 1.7 mM MgCl2; (Mudgett & Staskawicz 1999; Barrett *et al.* 2011)) and grew cultures overnight in a shaking incubator (250 rpm) at 28 C. MM at pH 5.6 induces expression of the type-III secretion system (T3SS) in a diversity of *Pseudomonas* spp. (Huynh *et al.* 1989), in contrast to KB, which results in negligible T3SS expression. T3SS expression was important for maximizing the potential relevance of our *in vitro* assay environments to those of plants, in which T3SS expression is expected. Thus, MM was used for this and all other culture experiments in this study.

Overnight cultures of each strain in MM were spun down for 3 m at 3000 $\times$ g and the supernatant replaced with 500 $\mu$L fresh MM. The density of each culture was adjusted to $OD_{600} = 0.2$ prior to 1:100 dilution into a total of 180 $\mu$L MM inside the wells of sterile polystyrene 96-well plates (Falcon). Each 96-well plate was covered with optically clear, gas-permeable plastic tape (Sigma `Z380059`) and incubated for 60 hr in a BioTek 600 plate reader in which $OD_{600}$ measurements were taken every 5 min with continuous random orbital shaking. Identical growth assays were performed on separate days in duplicate.

##### Estimating traits related to growth

We used **R** package `grofit` (Kahm *et al.* 2010) to fit smoothed functions to the bacterial growth data. Curve fits generated using logistic, Richards, Gompertz, or modified Gompertz equations

[*] Code and data available at `https://github.com/phumph/competitive_hierarchies`.

failed to produce estimates with $r \geq 0.5$ and we therefore used a non-parametric locally-weighted smoothing function to estimate the following growth curve parameters: maximum growth rate $r_m$, lag phase $l$, and maximum yield $K$. Lag phase represents the length of time (min) prior to initiation of exponential growth, while maximum yield is the maximum $OD_{600}$ attained during 60 h of growth. Curves for long lag-phased strains never leveled off (Fig. S1, e.g., strain `17A`); in these cases, $K$ was set as the final $OD_{600}$. When growth trajectories exhibited multiple exponential phases (diauxic shifts), $r_m$ was estimated during the initial exponential phase (e.g., strain `20A`; Fig. S1).

#### Pairwise competition assays

In 100 mm diameter Petri dishes, we overlaid 4 mL of 0.5% (w/v) soft agar inoculated with a resident strain overtop a sterile base layer of growth medium (minimal medium + 1% w/v agar). The top agar overlay for each plate contained a suspension of a single resident strain inoculated at $5 \times 10^5 \text{ml}^{-1}$ while soft agar was still molten ~42 C. Once cooled, suspensions of each of the 40 invader strains were then spotted at the same concentration in 4 $\mu$L aliquots spaced every 0.5 cm in parallel rows using an 8-channel pipettor. Resident and invader strain suspensions were made from exponential phase cultures in MM 3 mL with shaking at 28 C. Plates were incubated at 28 C face up for 12 h and then face down incubation for an addition 10 days. Megacolony spots were scored by hand for growth after days 1, 3, 5, 8, and 10. Data used for the following analyses are from day 10, by which time all interactions dynamics had leveled off.

### Supplemental Figures

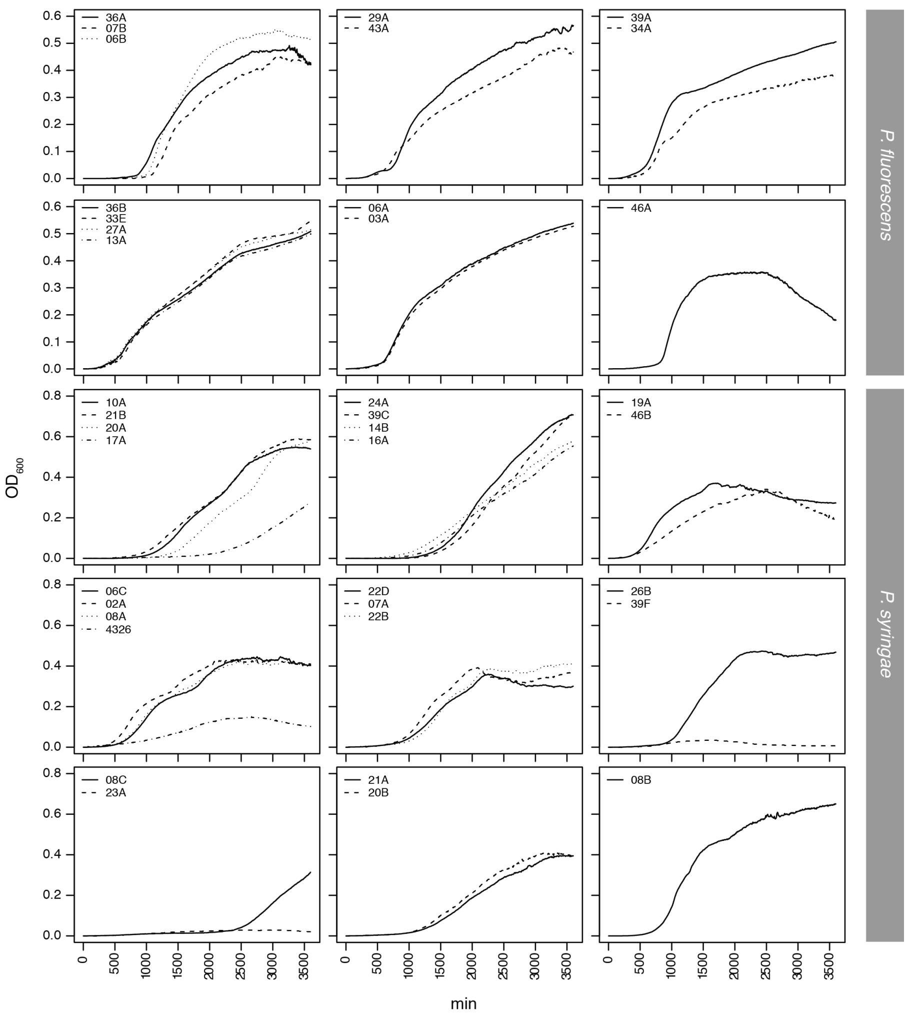


Figure S1: **Growth cycle profiles of 40 *Pseudomonas* spp. strains measured individually in minimal media *in vitro*.** Overlain individual growth curves are shown grouped by nearest phylogenetic neighboring strains (see Fig. 1, main text). Strain `08B` is sister to *P. syringae* clade. `4326` = laboratory strain *P. syringae* pv. maculicola str. `ES4326`.

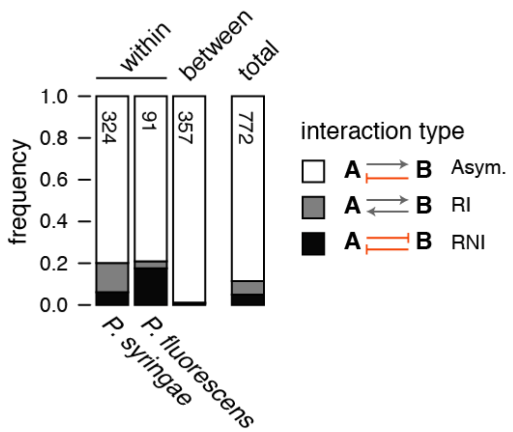


Figure S2: **Distribution of interaction outcomes within and between *Pseudomonas* clades**. Accompanying outcome counts and statistical results are displayed in Tables S1 and S2, respectively. Total strain pairings = 772 between 40 strains, with the 20 self-interactions and 8 no-data interactions removed. `RI` = reciprocal invasibility; `RNI` = reciprocal non-invasibility; `Asymm.` = asymmetric dominance.

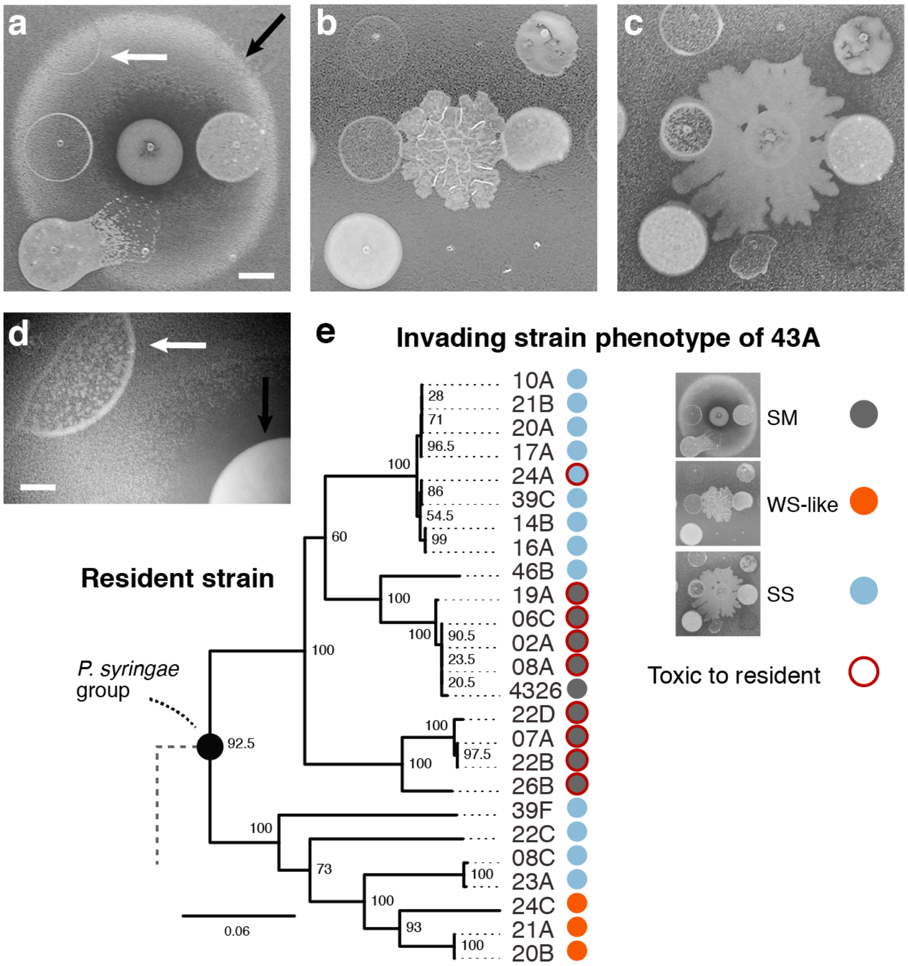


Figure S3: **Morphological variation observed during pairwise inhibition assays reflects strain-specific induction of motility and inhibition phenotypes in *P. fluorescens*. a-c**. Photos of interaction plates against *P. syringae* resident strains against which *P. fluorescens* strain 43A adopted distinct megacolony morphologies (white arrows). Black arrows denote invading strains producing tox+ halos against the resident strains. **d-f**. close-ups of smooth morph (SM), wrinkly spreader-like (WS-like), and the smooth spreader (SS) morphs from above. **g**. Close-up of putative growth facilitation of non-focal strain via secretion of effectors from 43A, arising from either cell cycle modification by secreted regulators, or competitive release via killing of resident strain. H. Phylogenetic specificity of morphological induction and its relationship with toxicity for strain 43A. Phylogenetic tree from Fig. 1 (main text). Scale bars in **a–c** = 1 cm; **d–f** = 0.50 cm; **g** = 0.25 cm.

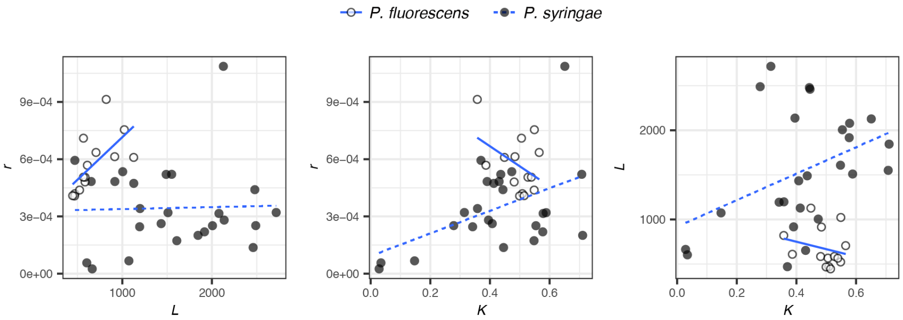


Figure S4: **Life history trait correlations exhibited by *P. syringae* and *P. fluorescens* strains.** Slopes of clade-wise linear regression are over-plotted in each panel for illustration purposes only regardless of $p$ value of correlation coefficient (see Fig. 1c, main text).

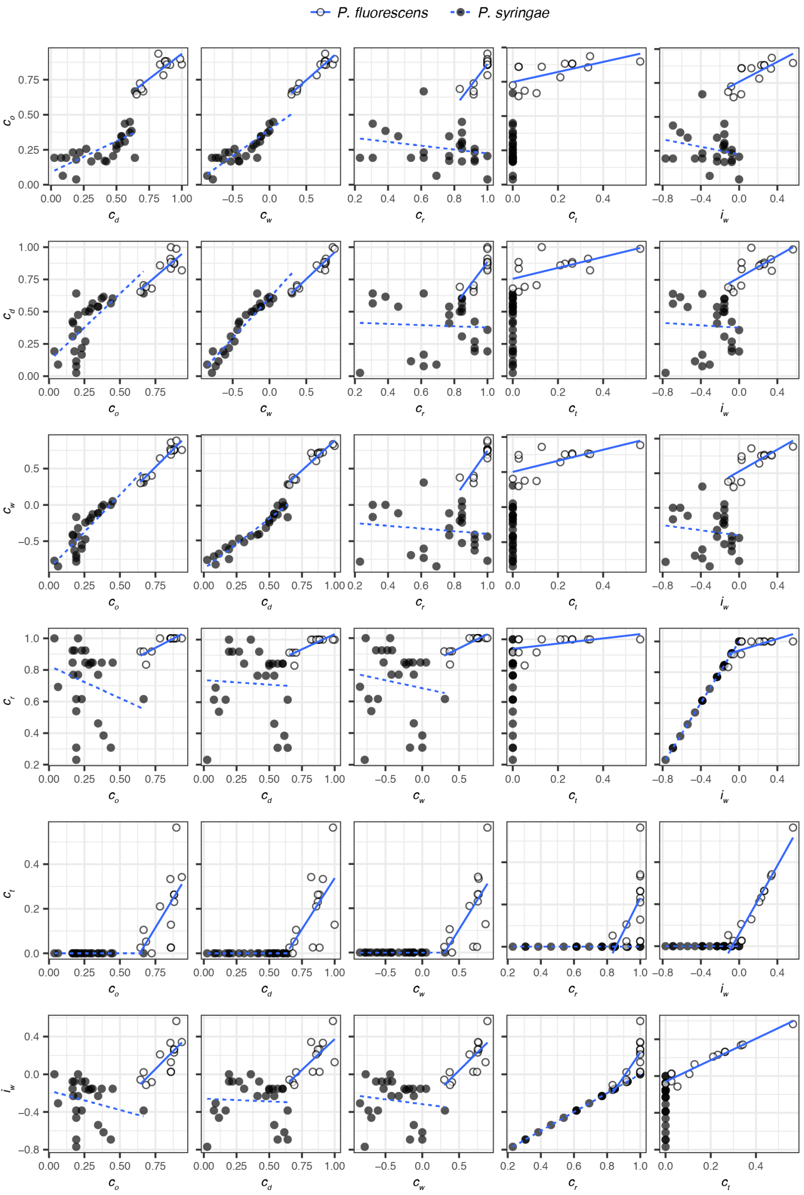


Figure S5: **Correlations among competitive traits exhibited by *P. syringae* and *P. fluorescens* strains.** Note that $C_t$ for *P. syringae* are all 0 because none produced detectable inhibitors scored in our pairwise soft-agar competition assays (see Extended Data Fig. 1, main text). Slopes of clade-wise linear regression are over-plotted in each panel for illustration purposes only regardless of $p$ value of correlation coefficient (see Fig. 1c, main text).

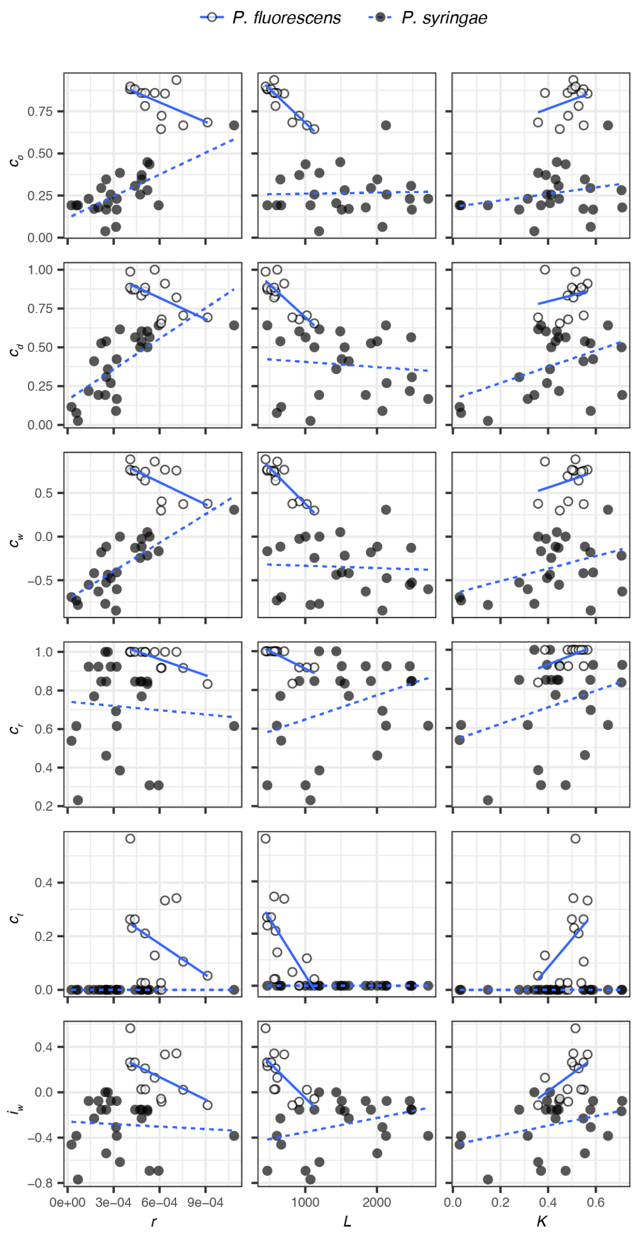


Figure S6: **Correlations between competitive traits and life-history traits exhibited by *P. syringae* and *P. fluorescens* strains.** Slopes of clade-wise linear regression are over-plotted in each panel for illustration purposes only regardless of $p$ value of correlation coefficient (see Fig. 1c, main text).

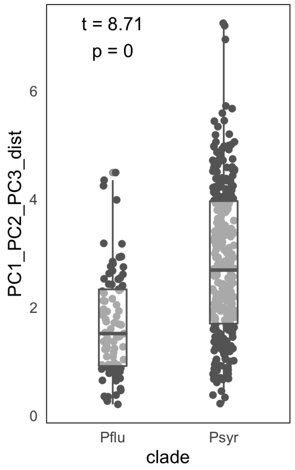


Figure S7: **Multivariate trait dispersion within *P. syringae* and *P. fluorescens* clades**. Pairwise Euclidean distances along trait PCs 1-3 are greater for *P. syringae* strains than among *P. fluorescence* strains (unequal variance two-sample Welch's $t$-test, $t = 8.71; p < 10^{-5}$).

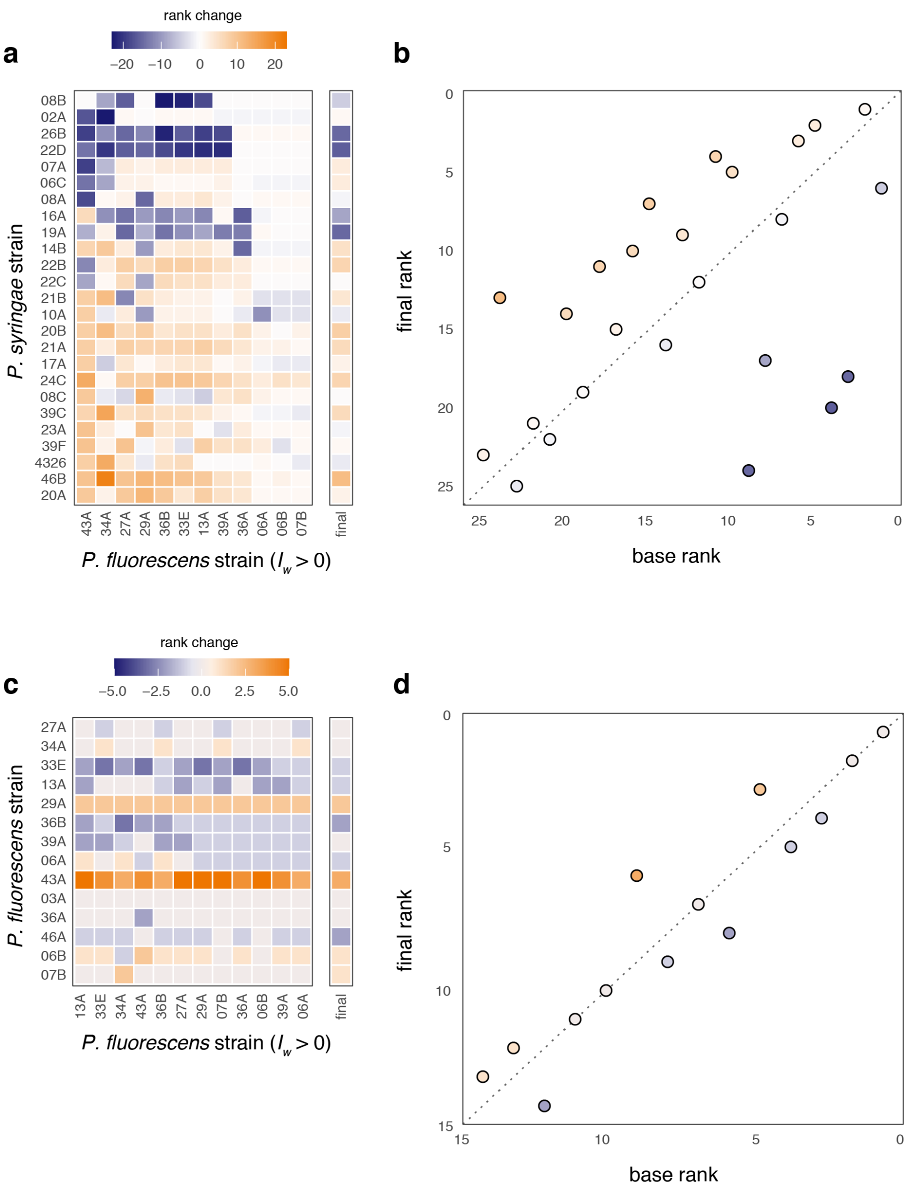


Figure S8: **Strain-wise and overall rank differences with and without indirect interactions from secretion-producing *P. fluorescens* strains. a-b.** Indirect effects on *P. syringae* reduce fitness most for strains with high base rank and facilitate weaker strains. Community-wide effects of indirect facilitation is to weaken fitness hierarchies among *P. syringae* strains (base rank versus final rank rank correlation = 0.52). **c-d.** In *P. fluorescens*, fitness ranks are far less affected by intra-clade indirect effects than in *P. syringae* (base rank versus final rank rank correlation = 0.94). Strains along the $y$ axis are plotted from high (top) to low (bottom) baseline fitness ($C_w$), while *P. fluorescens* strains are plotted from most facilitating (left) to least facilitating (right) along the $x$ axis.

### Supplemental Tables

Table S1: Summary statistics of life history and competitive trait distributions.

| trait | *P. fluorescens* | *P. syringae* |
|---|---|---|
| $\mu(\sigma)$ | | |
| $r$ | 0.57 (0.15) | 0.34 (0.22) |
| $L$ | 672.13 (216.43) | 1549.86 (656.52) |
| $K$ | 0.49 (0.06) | 0.42 (0.18) |
| $i_w$ | 0.15 (0.19) | -0.28 (0.22) |
| $c_w$ | 0.65 (0.2) | -0.35 (0.3) |
| $c_t$ | 0.18 (0.16) | 0 (0) |
| $c_r$ | 0.97 (0.05) | 0.72 (0.22) |
| $c_o$ | 0.82 (0.1) | 0.26 (0.13) |
| $c_d$ | 0.83 (0.11) | 0.39 (0.2) |
| **[min; max]** | | |
| $r$ | [0.41; 0.91] | [0.03; 1.09] |
| $L$ | [446.1; 1126.92] | [470.52; 2716.58] |
| $K$ | [0.36; 0.57] | [0.03; 0.71] |
| $i_w$ | [-0.11; 0.56] | [-0.77; 0] |
| $c_w$ | [0.3; 0.88] | [-0.85; 0.31] |
| $c_t$ | [0; 0.56] | [0; 0] |
| $c_r$ | [0.83; 1] | [0.23; 1] |
| $c_o$ | [0.64; 0.94] | [0.04; 0.67] |
| $c_d$ | [0.65; 1] | [0.03; 0.64] |

[a] $r$ displayed as $\times 1000$

Table S2: Distribution of outcomes by pairing type.

| predictor | ASYM | RI | RNI |
|---|---|---|---|
| **Counts** | | | |
| Between | 353 | 2 | 2 |
| Within (*P. fluorescens*) | 72 | 16 | 3 |
| Within (*P. syringae*) | 259 | 20 | 45 |
| **Frequencies** | | | |
| Between | 0.989 | 0.006 | 0.006 |
| Within (*P. fluorescens*) | 0.791 | 0.176 | 0.033 |
| Within (*P. syringae*) | 0.799 | 0.062 | 0.139 |

[a] RNI = Reciprocal non-invasion
[b] RI = Reciprocal invasion
[c] ASYM = Asymmetric dominance

Table S3: Multinomial model coefficients

| outcome | term | coefficient[a] | std.err | $z$ | $p-value$ |
|---|---|---|---|---|---|
| RI[b] | intercept | 0.01 | 0.71 | -7.30 | $< 10^{-5}$ |
| RI | *P. fluorescens* | 39.22 | 0.76 | 4.82 | $< 10^{-5}$ |
| RI | *P. syringae* | 13.63 | 0.75 | 3.50 | 0.0005 |
| RNI[c] | intercept | 0.01 | 0.71 | -7.30 | $< 10^{-5}$ |
| RNI | *P. fluorescens* | 7.36 | 0.92 | 2.16 | 0.031 |
| RNI | *P. syringae* | 30.67 | 0.73 | 4.71 | $< 10^{-5}$ |

[b] RI = Reciprocal invasion
[a] RNI = Reciprocal non-invasion
[c] coefficients = odds

Table S4: Linear regression of trait distance versus phylogenetic distance by clade.

| term | estimate | std.error | statistic | p.value |
|---|---|---|---|---|
| intercept | 1.03 | 0.17 | 6.22 | $< 10^{-5}$ |
| $D_g$ | 0.09 | 0.01 | 6.46 | $< 10^{-5}$ |
| clade (*P. syringae*) | 0.88 | 0.15 | 5.76 | $< 10^{-5}$ |
